# Tertiary structure of a Nif11-like leader peptide mediates protein-protein interactions with a RiPP biosynthetic enzyme

**DOI:** 10.64898/2026.10.04.756588

**Authors:** Mujeeb A. Wakeel, FNU Vidya, Collin E. Merrick, Lauren N. Mullininx, Estella F. Yee, John S. Harwood, Elizabeth I. Parkinson, Andrew C. McShan, Vinayak Agarwal

## Abstract

Ribosomally synthesized and post-translationally modified peptides (RiPPs) are produced from precursor peptides that typically contain a short, unstructured N-terminal leader region that engages RiPP biosynthetic enzymes through peptide-protein contacts. A distinct subclass of leader peptides, the nitrogen-fixing protein 11-like leader peptides (Nif11-LPs), are unusually long and conspicuously enriched in cyanobacteria. No atomic structures of Nif11-LPs have been reported. Here, we report the solution structure of MppE^leader^, a Nif11-LP from the marine cyanobacterium *Moorena producens* PAL. MppE^leader^ adopts a bipartite architecture with a rigid N-terminal tetrahelical bundle stabilized around a hydrophobic nucleus, followed by a dynamic, largely disordered C-terminal region containing a conserved L(X)₄L motif. We show that the α-ketoglutarate/non-heme iron enzyme MppG dihydroxylates the macrocyclized MppE^core^ peptide in a strictly leader-dependent manner. NMR titrations and structure-guided mutagenesis demonstrate that both the tetrahelical hydrophobic nucleus and the L(X)₄L motif of MppE make direct, functionally required contacts with MppG, including its RiPP recognition element (RRE) domain. An AlphaFold3-guided structural model rationalizes these contacts. Together, these findings establish a tertiary structure-dependent protein-protein interaction paradigm for RiPP core peptide modification, distinguishing Nif11-LP recognition from canonical short-leader mechanisms and expanding the understanding of substrate recognition in RiPP biosynthesis.

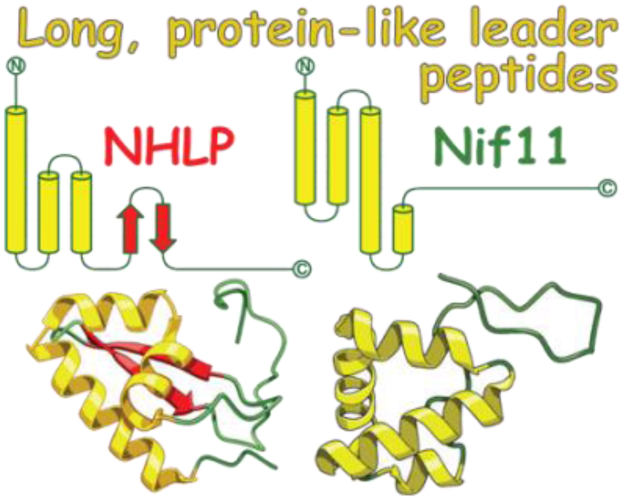

## Introduction

The biosynthesis of ribosomally synthesized and post-translationally modified peptides (RiPPs) involves tailoring of a precursor peptide by RiPP biosynthetic enzymes.^1^ Genes encoding the precursor peptide and the biosynthetic enzyme are typically present in genetic neighborhoods called biosynthetic gene clusters (BGCs). Genome mining has established the ubiquity of RiPP encoding BGCs across all forms of life.^2^ The discovery of RiPPs and RiPP encoding BGCs is motivated by the established pharmaceutical utility of peptidic molecules as well as their demonstrated roles in intra- and inter-organismal biology.^3,4^

Chemical modifications underlying RiPP biosynthesis expands the chemical space explored by ribosomally synthesized peptides.^5,6^ These modifications involve tailoring the peptide backbone as well as the amino acid side chains.^1^ Among others, peptide backbone modifications include thioamidation,^7^ Cα epimerization,^8^ backbone heterocycle formation,^9,10^ and installation of β-amino acids.^11^ The inventory of side chain modifications is extraordinarily vast. A convergent theme in RiPP biosynthesis is macrocyclization—affected using various C–C and C–heteroatom bond forming strategies. Macrocyclization rigidifies the RiPP structure, lending it proteolytic stability, and facilitates transport through membranes.^12^

Divested from RiPP biosynthesis, chemical modification of peptides and proteins is also involved in primary biological processes, such as signal transduction, epigenetic regulation, and production of peptidic hormones, toxins, and venoms. There is overlap among the post-translational modifications (PTMs) involved in RiPP biosynthesis, and PTMs in primary biology. Examples include thioamidation, thionylation, methylation, glycosylation, and disulfide bond formation.^13–18^ While the PTMs in themselves may be chemically similar, molecular recognition of a precursor peptide by RiPP biosynthetic enzymes can be vastly different as compared to recognition of large proteins that possess tertiary and quaternary structures by PTM-installing enzymes in primary biology. RiPP precursor peptides are usually divided between an N-terminal leader peptide region and a C-terminal core region. Often, the leader mediates interaction between the precursor peptide and the RiPP biosynthetic enzyme but is itself not modified. Rather, PTMs are affected upon the core, which is subsequently cleaved from the unmodified leader to furnish the mature RiPP.^1^

The canonical RiPP leader peptides are short, usually fewer than 50 amino acids, and largely devoid of folded structure in isolation.^19^ Several structural descriptions for leader peptide binding to RiPP biosynthetic enzymes are now available.^20–24^ These interactions *typically* involve hydrophobic side chains on the leader peptide binding to a complementary register of hydrophobic grooves on the RiPP biosynthetic enzyme.^20,21^ Interactions can also be mediated by the RiPP recognition element (RRE) domains that are embedded within or associated with RiPP biosynthetic enzymes.^25^ In RRE-mediated interactions, the unstructured leader peptide adopts an isolated β-strand that extends the anti-parallel β-sheet of the RRE domain.^22,23^

In contrast to the canonical short leader peptides, two families of RiPP precursor leader peptides have been described that are significantly longer in length (>75 amino acids). These are the nitrile hydratase-like leader peptides (NHLPs) and the nitrogen fixing protein 11-like leader peptides (Nif11-LPs.^26^ Experimentally determined structures reveal that NHLPs possess protein-like rigid tertiary folds followed by intrinsically disordered peptide-like extensions.^27,28^ The tertiary structures of the N-terminal structured domains of NHLPs have been demonstrated to be involved in intermolecular interactions with RiPP biosynthetic enzymes.^28,29^ Thus, rather than the peptide/protein interaction regime used by the short, canonical RiPP leader peptides, the interaction of NHLPs with RiPP biosynthetic enzymes follows a fundamentally different protein/protein interaction paradigm wherein tertiary structural interactions may be involved. Akin to NHLPs, the Nif11-LPs are atypically long and predicted to have well-defined tertiary structures. However, atomic structures and RiPP biosynthetic enzyme binding modes for Nif11-LPs remain undeciphered.

In this study, a combination of solution-state nuclear magnetic resonance (NMR) spectroscopy and small angle X-ray scattering (SAXS) was used to determine the three-dimensional structure of MppE^leader^, a Nif11-LP from the marine cyanobacterium *Moorena producens* PAL. Solution NMR and molecular dynamics (MD) simulations highlight differences in the dynamic behavior of the N-terminal rigid and C-terminal intrinsically disordered regions. Guided by NMR titrations, computational modeling, and mutagenesis, we biochemically validate the protein/protein interactions that guide core peptide modification by an α-ketoglutarate-dependent mononuclear non-heme iron hydroxylase (NHFe/α-KG hydroxylase). These findings illuminate a new modality for substrate recognition in RiPP biosynthesis. Here, the protein/protein interactions rely on the tertiary structure of a Nif11-LPs and a sequence motif within the Nif11-LPs that interacts with the RRE domain within the NHFe/α-KG hydroxylase.

## Results and Discussion

### Cyanobacteria are prolific sources of Nif11-LP containing RiPP precursor peptides

The phylogenetic distribution of NHLPs has been described before to reveal their spread across different bacterial phyla.^26,30,31^ In contrast, mining the available sequence space reveals that Nif11-LPs are unusually enriched in cyanobacteria. Cyanobacteria are exceptionally rich sources for RiPP precursor peptides possessing Nif11-LPs with nearly two-third of all Nif11-LP sequences derived from the phylum Cyanobacteriota (Fig. 1). Curiously, multiple RiPP precursor peptides possessing Nif11-LPs are spread across cyanobacterial genomes oblivious to the traditional gene neighborhood definition of natural product BGCs.^32,33^ Correspondingly, the associated RiPP biosynthetic enzymes that modify these precursor peptides have proven to be substrate promiscuous biotechnology tools.^34^ Given their enrichment in cyanobacterial genomes, it is not surprising that RiPPs generated from precursor peptides possessing Nif11-LPs play fundamental roles in cyanobacterial ecology.^35,36^

**Fig. 1.**
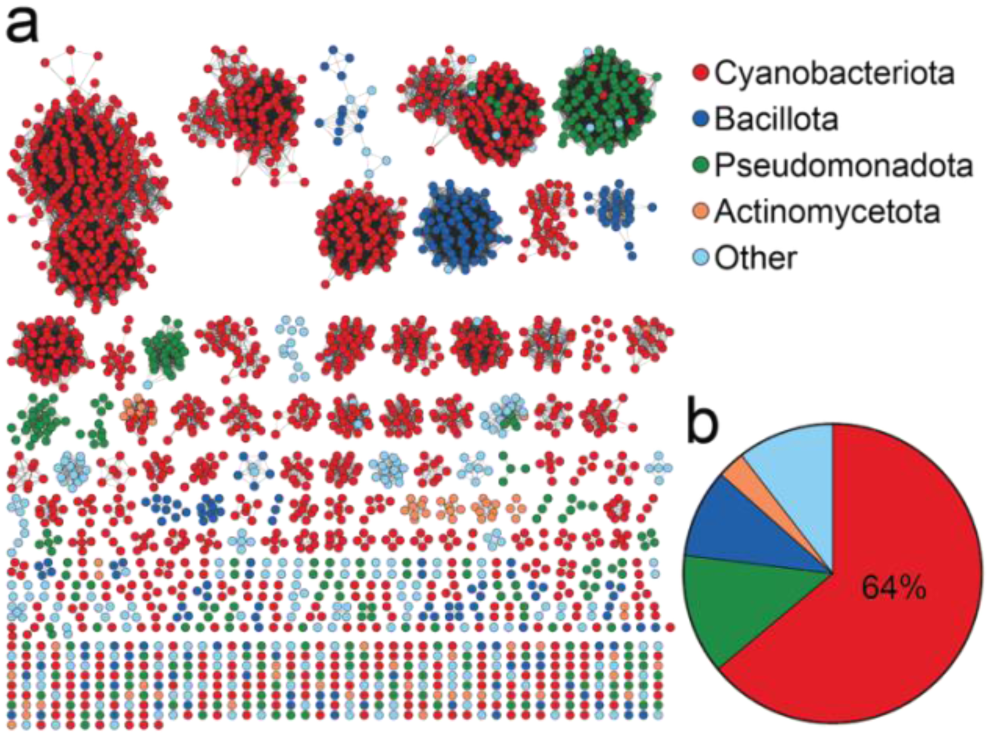
Nif11-LP sequence space. **a** Sequence similarity network (SSN) with nodes colored according to selected bacterial phyla. Cyanobacterial nodes are colored red. **b** Of the 2,395 nodes represented in the SSN, 64% are derived from Cyanobacteriota thus demonstrating that this phylum as the dominant source of RiPP precursor peptides possessing Nif11-LPs.

### Activity of a Nif11-LP dependent peptide hydroxylase

We have recently reported the discovery of a cryptic RiPP-encoding *mpp* BGC from the filamentous marine cyanobacterium *Moorena producens* PAL (Fig. 2a).^37,38^ Within the *mpp* BGC were identified genes encoding a lanthionine synthetase MppM, a flavin-dependent brominase MppI, prenyltransferase MppC, and a RiPP precursor peptide MppE.^38^ The 77 amino acid-long leader sequence for precursor peptide MppE—henceforth referred to as MppE^leader^—belongs to the Nif11-LP family. The MppE^leader^ is appended to a heptapeptidic MppE^core^ with the sequence ACWRWSG (Table S1). While the final bioactive RiPP(s) generated by the *mpp* BGC remain cryptic, transformations affected upon MppE by MppM, MppI, and MppC have been reported (Fig. 2b).^38,39^ However, the functions of other enzymes encoded in the *mpp* BGC remain unknown. The focus of this report is the activity of the enzyme MppG encoded in the *mpp* BGC and its interactions with the Nif11-LP MppE^leader^.

**Fig. 2.**
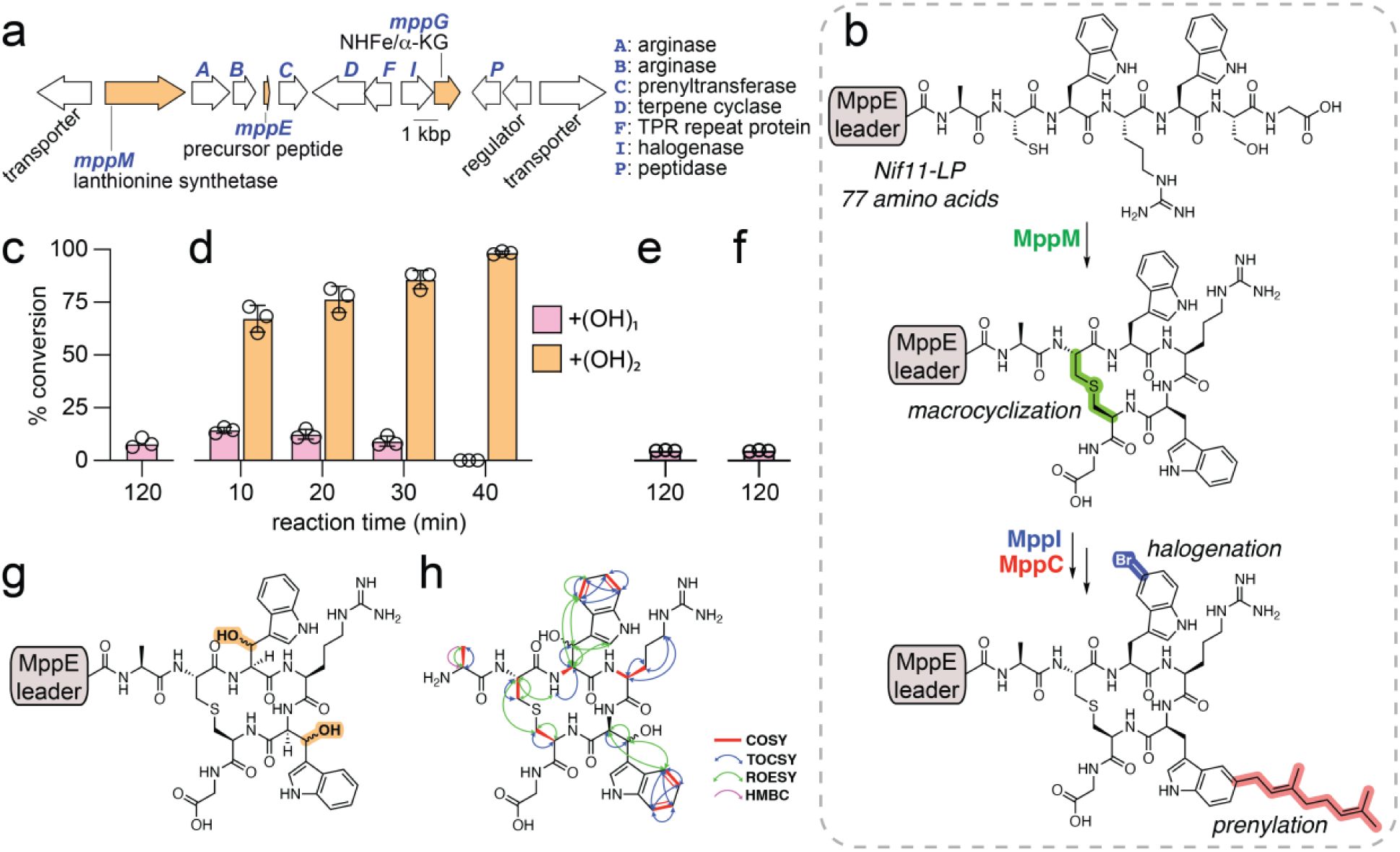
NHFe/α-KG MppG modifies macrocyclized MppE. **a** The *mpp* BGC showing the predicted or experimentally established activities of key genes. Genes studied in this work are highlighted orange. **b** Scheme showing post-translational tailoring of the MppE precursor peptide by enzymes with established activities. Macrocyclization by MppM precedes halogenation by MppI and prenylation by MppC. **c** Extent of turnover to monohydroxylated product in 120 min when unmodified linear MppE was provided as a substrate to MppG. **d** Time-dependent turnover of MppM-modified, macrocyclized MppE to mono- and dihydroxylated products by MppG. Note the difference in reaction times between panels c and d. **e** Turnover to monohydroxylated product when the macrocyclized MppE^core^ alone, without the MppE^leader^, was provided as a substrate to MppG. No dihydroxylated product formation was observed. **f** Turnover to monohydroxylated product by MppG when the excised MppE^leader^ was provided *in trans* to the macrocyclized MppE^core^ as substrate. No dihydroxylated product formation was observed. **g** Structure of the dihydroxylated product furnished by MppG. Configuration of the secondary alcohol remains undefined. **h** Key NMR correlations for the dihydroxylated MppE^core^ product furnished by MppG. Note that the MppE^leader^ was proteolytically removed to yield the macrocyclized and dihydroxylated MppE^core^ prior to spectroscopic characterization.

Based on sequence similarity, MppG was predicted to be an NHFe/α-KG oxygenase. An AlphaFold 3-predicted model of MppG revealed a Fe-ion coordinating His/Asp triad (pTM=0.76, ipTM=0.98, Fig. S1). Members of the NHFe/α-KG family of enzymes catalyze myriad chemically challenging transformations.^40^ In the context of RiPP biosynthesis, NHFe/α-KG enzymes have been reported to catalyze hydroxylation of amino acid side chains.^41–46^ First, we explored the activity of MppG against an unmodified MppE precursor peptide. Even with long reaction time, only modest hydroxylation (<10% conversion) of the unmodified, linear MppE precursor peptide was observed in aerobic *in vitro* reactions employing purified MppG along with α-KG, Fe(II), and ascorbate (Fig. 2c, Fig. S2–S3). The hydroxylated product was highly labile; under acidic solvent conditions employed for liquid chromatography/mass spectrometry (LC/MS) experiments, we observed concomitant appearance of ions corresponding to a dehydrated species (−H_2_O, Fig. S4). Next, we tested macrocyclized MppE as a substrate wherein a lanthionine ring has been pre-installed by MppM employing the MppE-Cys79 and MppE-Ser83 side chains (Fig. 2b).^38^ Rapid, time-dependent hydroxylation of the macrocyclized MppE^core^ was observed to be catalyzed by MppG (Fig. 2d). Eliminating MppG or α-KG from the reaction abolished product formation while the lack of addition of Fe(II) or ascorbate reduced product formation; these observations are in line with the established enzymology of NHFe/α-KG family of enzymes (Fig. S5).^47^ Even with reaction times of 10 min, we observed production of a dihydroxylated product (+(OH)_2_; Fig. 2d, Fig. S6). With increasing reaction times, the abundance of the monohydroxylated product decreased with the dihydroxylated product dominating (Fig. 2d). These observations allow us to posit that the physiological substrate of MppG is the MppM-modified macrocyclic MppE. The substrate preference of MppG conforms to previous descriptions of the activities of the halogenase MppI and prenyltransferase MppC that also preferentially modify macrocyclized MppE^core^ as compared to the linear MppE^core^. Furthermore, MppG is a processive enzyme that affects two distinct hydroxylation events upon the macrocyclized MppE^core^. The mono- and dihydroxylated products were highly labile and dehydrated products were detected.

Hydroxylation in RiPP biosynthesis by NHFe/α-KG family of enzymes can proceed in a leader-dependent, or in a leader independent fashion.^42,43^ To query the leader dependence on the hydroxylation activity of MppG, the macrocyclized MppE^core^ was excised from the MppE^leader^ using the LahT150 protease and provided as a substrate to MppG.^21^ Even with extended reaction time of 120 min, minimal turnover of the MppE^core^ in the absence of the MppE^leader^ to monohydroxylated product was observed; no dihydroxylated product formation was detected, at all (Fig. 2e, Fig. S7). When the excised MppE^leader^ was provided *in trans* along with the macrocyclized MppE^core^, as before, only minimal monohydroxylated product formation was detected (Fig. 2f, Fig. S8). These observations indicate that dihydroxylation of the macrocyclic MppE^core^ by MppG is leader dependent such that MppG would make intimate contact with the Nif11-LP MppE^leader^.

To establish the sites of dihydroxylation by MppG, the dihydroxylated product was produced in preparative amounts, excised from MppE^leader^ using trypsin and Glu-C proteases, and chromatographically purified. Comprehensive solution-state NMR analyses established that the β-positions of both Trp residues in the macrocyclized MppE^core^ were hydroxylated by MppG (Table S2, Fig. 2g–2h, Fig. S9–S15).

Hydroxylation at the Trp-Cβ positions is in line with the observation of rapid dehydration under acidic conditions wherein protonation of the alcohol would be accompanied by dehydration leading to C*a*-Cβ unsaturation in conjugation with the Trp indole (Fig. S16). The highly labile nature of the dihydroxylated product generated by MppG precluded experimental determination of the configuration of the secondary alcohol.

### Solution-state structure of Nif11-LP MppE^leader^

Observations described above establish that the hydroxylation activity of MppG is dependent on interactions with the Nif11-LP MppE^leader^. To query the interactions between the MppG and MppE^leader^, we first sought to determine the biophysical properties and atomic structure of MppE^leader^. Far-UV circular dichroism (CD) spectroscopy spectra of recombinant MppE^leader^ demonstrated characteristic negative dips at 208 and 222 nm, supporting a structure containing α-helices (Fig. 3a). Deconvolution of the CD spectra with BeStSel suggested that MppE^leader^ possesses primarily α-helical and disordered secondary structure elements (Fig. S17).^48^ Thermal denaturation experiments established the melting temperature (T_m_) of MppE^leader^ to be 58.9°C, indicative of stable tertiary structure (Fig. 3b).

**Fig. 3.**
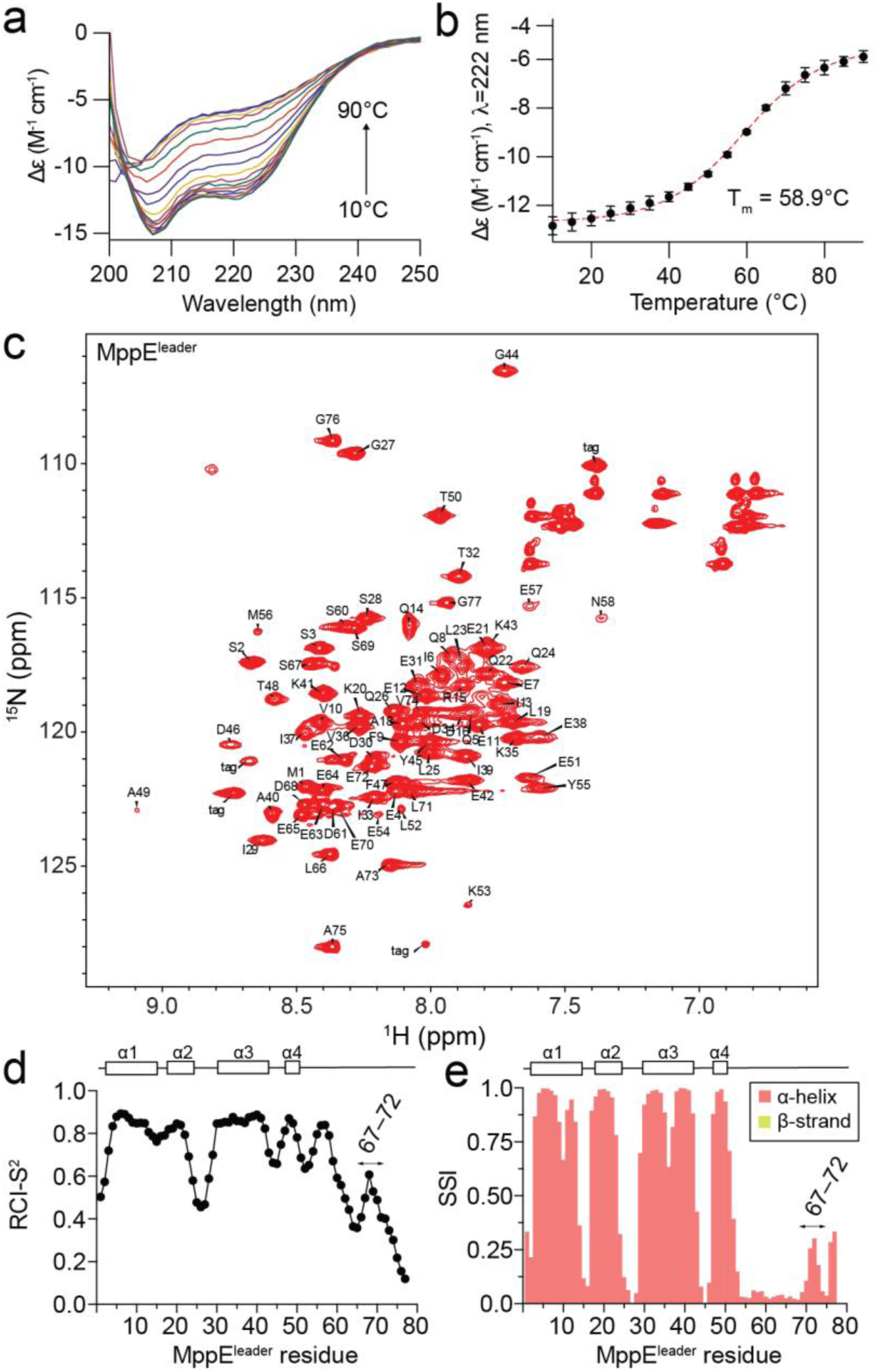
Biophysical characterization of the Nif11-LP MppE^leader^. **a** Far-UV CD spectra of MppE^leader^ measured from 10 to 90°C in 5°C increments. **b** CD thermal denaturation curve of MppE^leader^ acquired at 222 nm from 10 to 90°C. Melting temperature (T_m_) was determined from fitting with a Boltzmann sigmoidal equation (dotted red line). **c** 2D ^1^H-^15^N HSQC spectra of U-[^15^N,^13^C]-labeled MppE^leader^ acquired at a ^1^H field of 800 MHz at 25°C. Resonance assignments of NMR peaks corresponding to backbone amides are provided. NMR peaks corresponding to the remnants of the affinity purification tag are denoted “tag”. **d** TALOS-N derived RCI-S^2^ (black scatter, 0 and 1 signify disorder and rigid structure, respectively). **e** TALOS-N derived SSI (salmon = α helix, yellow = β-strand). RCI-S^2^ and SSI values are derived from assigned backbone chemical shifts. Residues 67–72 that possess slightly higher RCI-S^2^ and SSI values are indicated.

Next, uniformly ^15^N and ^13^C isotopically labeled (U-[^15^N,^13^C]) MppE^leader^ was prepared. We obtained 97% chemical shift assignment of backbone H_N_, NH, C_α_, C_β_, and C’ atoms using sequential assignment techniques based on three-dimensional triple-resonance NMR experiments (Fig. 3c).^49^ The random coil index order parameter (RCI-S^2^) and the secondary structure index (SSI) were calculated from chemical shifts using TALOS-N.^50^ RCI-S^2^ values suggested that the N-terminus of MppE^leader^ (residues 1–52) was highly ordered with increasing disorder towards the C-terminus (residues 53–77) (Fig. 3d). SSI supported the presence of α-helical character at the N-terminus of MppE^leader^ followed by an intrinsically disordered region towards the C-terminus (Fig. 3e). SSI and RCI-S^2^ plots both provide hints of a transiently ordered short α-helix in the MppE^leader^ at the C-terminus (residues 67–72) (Fig. 3d–3e). Together, these data support that recombinant MppE^leader^ behaves as a folded protein in solution that contains both rigid α-helical and intrinsically disordered characteristics.

We combined the chemical shifts, dihedral angle restraints from TALOS-N, and amide-to-amide NOE-based distance constraints within CS-Rosetta to solve the solution-state NMR structure of the MppE^leader^, which represents the first reported atomic structure of a Nif11-LP (Fig. 4a, Table S3). The MppE^leader^ N-terminal region packs into a four α-helix bundle spanning residues 2–15, 17–25, and 37–43, and 48–51. Among the ten energy minimized structures, this tetrahelical bundle adopted a conserved configuration (Fig. 4b).

**Fig. 4.**
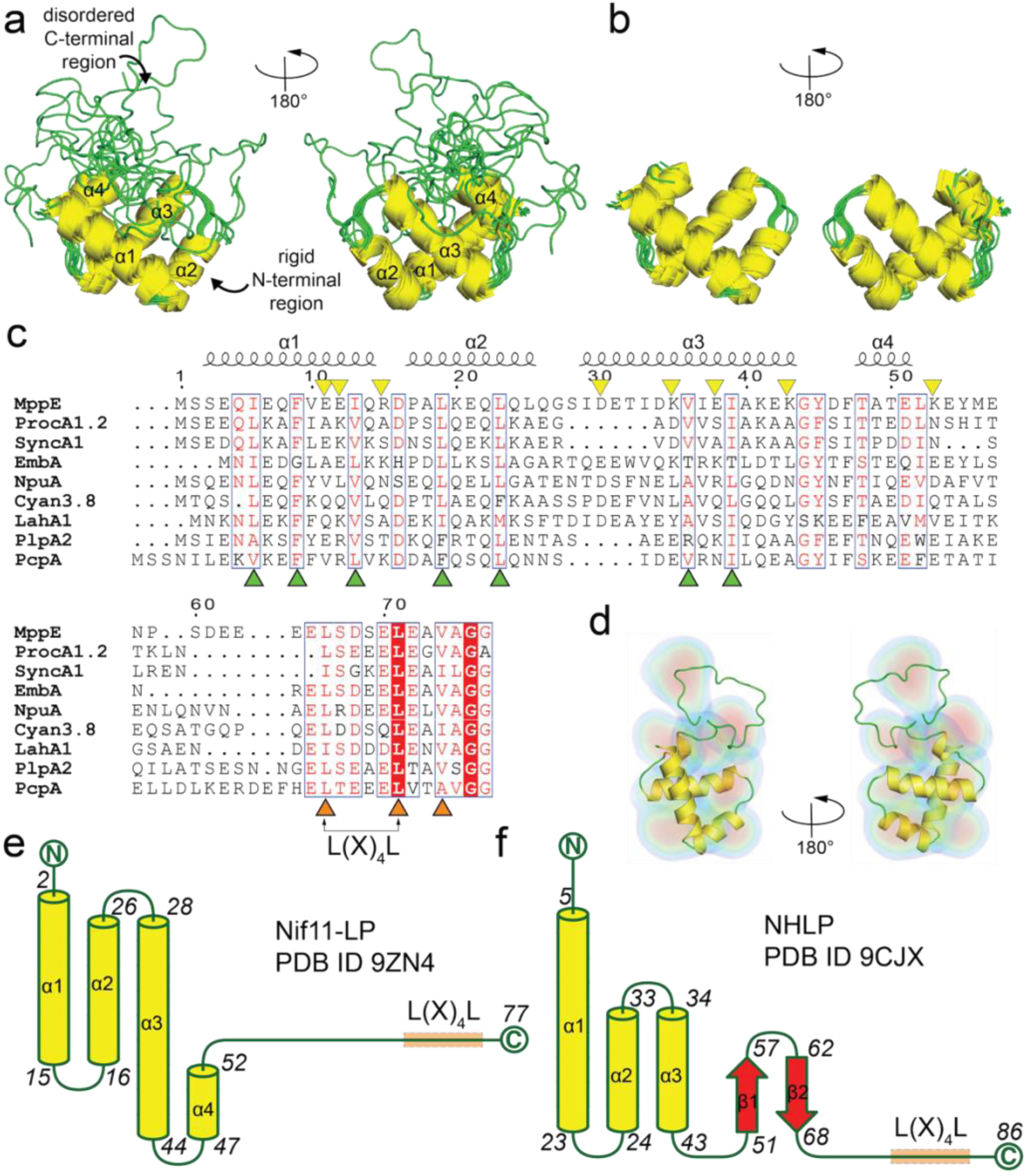
Solution NMR structure of the Nif11-LP MppE^leader^. **a** Two views for ten lowest energy solution structures of MppE^leader^ in cartoon representation. The α-helices are colored yellow, and the loops are colored green. **b** Two views in cartoon representation for ten lowest energy solution structures of rigid N-terminal domain of MppE^leader^ that comprises of a bundle of four α-helices. **c** Sequence alignment of Nif11-LPs. The green triangles denote residues that construct the hydrophobic nucleus in MppE^leader^, and the yellow triangles denote residues that are involved in salt bridge interactions observed in the NMR structure of the MppE^leader^. Toward the C-terminus of the MppE^leader^, orange triangles denote the −4, −7, and −12 residues that direct the activity of C39 peptidases; the −7 and −12 Leu residues construct the L(X)_4_L motif. **d** NMR structure of MppE^leader^ (red helices, green loops) overlaid on a DENSS-reconstructed electron density map calculated from SAXS experiments. Highest to lowest densities are rendered with a color ramp from red to blue using PyMOL v2.6.2. The best fitting NMR model was determined by FoXS and aligned to the DENSS map using ChimeraX. Refined DENSS map χ^2^ = 0.88. Secondary structure topology diagrams for **e** Nif11-LP MppE^leader^ (PDB ID 9ZN4) and **f** NHLP DprE2^leader^ (PDB ID 9CJX). Unlike NHLPs, Nif11-LPs do not possess any β-strands. Note that the L(X)_4_L motif is conserved among both NHLPs and Nif11-LPs and is positioned on the disordered region proximal to the C-terminus.

The structure of MppE^leader^ can be rationalized in context of sequence similarity to other Nif11-LPs (Fig. 4c). Reminiscent of the tetrahelical architecture of acyl carrier proteins,^51^ the MppE^leader^ structured region is stabilized around a hydrophobic nucleus composed of side chains from residues Ile6, Phe9, Val10, Ile13, Leu19, Leu23, Ile33, Val36, and Ile39 (Fig. S18). Most of these residues are conserved among Nif11-LPs (Fig. 4c; green triangles). This observation allows us to posit that burial of a hydrophobic nucleus by the tetrahelical structured domain would be a conserved feature among Nif11-LPs. Note that similar hydrophobic nuclei have also been observed in solution structures of NHLPs.^27,28^ Salt bridges between Glu11/Arg15, Glu12/Lys43, and Lys35/Glu38 side chains further rigidify the MppE^leader^ tetrahelical bundle. An additional salt bridge between the side chains of Asp30 and Lys53 can be discerned in some, but all not all, energy minimized structures. The salt bridge forming residues are not conserved among Nif11-LPs (Fig. 4c; yellow triangles). This could imply that the register of salt bridge residues could be variable among Nif11-LPs.

The C-terminal region of MppE^leader^, spanning residues 53–77, does not possess any well-defined secondary structural elements. It also does not appear to strongly associate with the tetrahelical bundle apart from the Met56 side chain that is positioned in toward the hydrophobic nucleus in some, but not all, energy minimized structures. Across the ten energy minimized structures, the four α-helices were found to be rigid, whereas the C-terminal region exhibits substantial conformational flexibility (Fig. 4a–4b). Electron density envelopes calculated using SAXS data supported the solution NMR-determined structures, indicating a globular monomer with a radius of gyration (R_g_) of 14.04 ± 0.03 Å (Fig. S19, Table S4). The dimensionless Kratky plot exhibited a semi-Gaussian profile, consistent with a protein containing an intrinsically disordered region, which is attributed to the conformational flexibility of the C-terminal region (Fig. S20–S21). Fitting of the NMR model into the SAXS-derived DENSS density map showed good agreement with a χ² value of 0.88 (Fig. 4d).

The bipartite structural organization of the Nif11-LP, where a rigid N-terminal region organized around a hydrophobic core is followed by a long unstructured C-terminal region, is reminiscent of the architecture of the NHLPs.^27,28^ The key structural distinction between NHLPs and Nif11-LPs is the presence of a two-strand antiparallel β-sheet present in NHLPs, which is substituted by the short, fourth α-helix in Nif11-related leader peptides (Fig. 4e–4f).

### Picosecond to nanosecond dynamics of MppE^leader^

Conformational dynamics of the NHLPs have been suggested to contribute to the interactions of the leader peptide with RiPP-modifying enzymes.^27,28^ To characterize the dynamic properties of the Nif11-LP MppE^leader^ on the picosecond-to-nanosecond timescale, we acquired HSQC-based R_1_ and R_2_ relaxation rates with ^15^N-{^1^H} heteronuclear NOE (HetNOE) values.^52^ R_1_, R_2_, and ^15^N-{^1^H} HetNOE data support the rigidity of the MppE^leader^ α1, α2, α3, and α4 helices (Fig. 5a). Taken together, the R_1_, R_2_, and HetNOE measurements are consistent with the solution NMR structure of MppE^leader^ and highlight the rigidity in the N-terminal region and flexibility within the C-terminal region. In addition, we also observed a small increase in R_1_ and ^15^N-{^1^H} HetNOE values for MppE^leader^ residues 67–72 (Fig. 5a), consistent with indications of transient rigidness implied by RCI-S^2^ and SSI values of these residues (Fig. 3d–3e). These observations are indicative of the possibility of a transiently populated ordered-state within the C-terminal tail of Nif11-LP MppE^leader^. Similar observations have been made previously for NHLPs.^28^ Of note, this region in Nif11-LPs and NHLPs comprises of the L(X)_4_L motif (Leu66–Leu71 for MppE) which has been shown to mediate interactions with RiPP biosynthetic enzymes.^27,29,38^

**Fig. 5.**
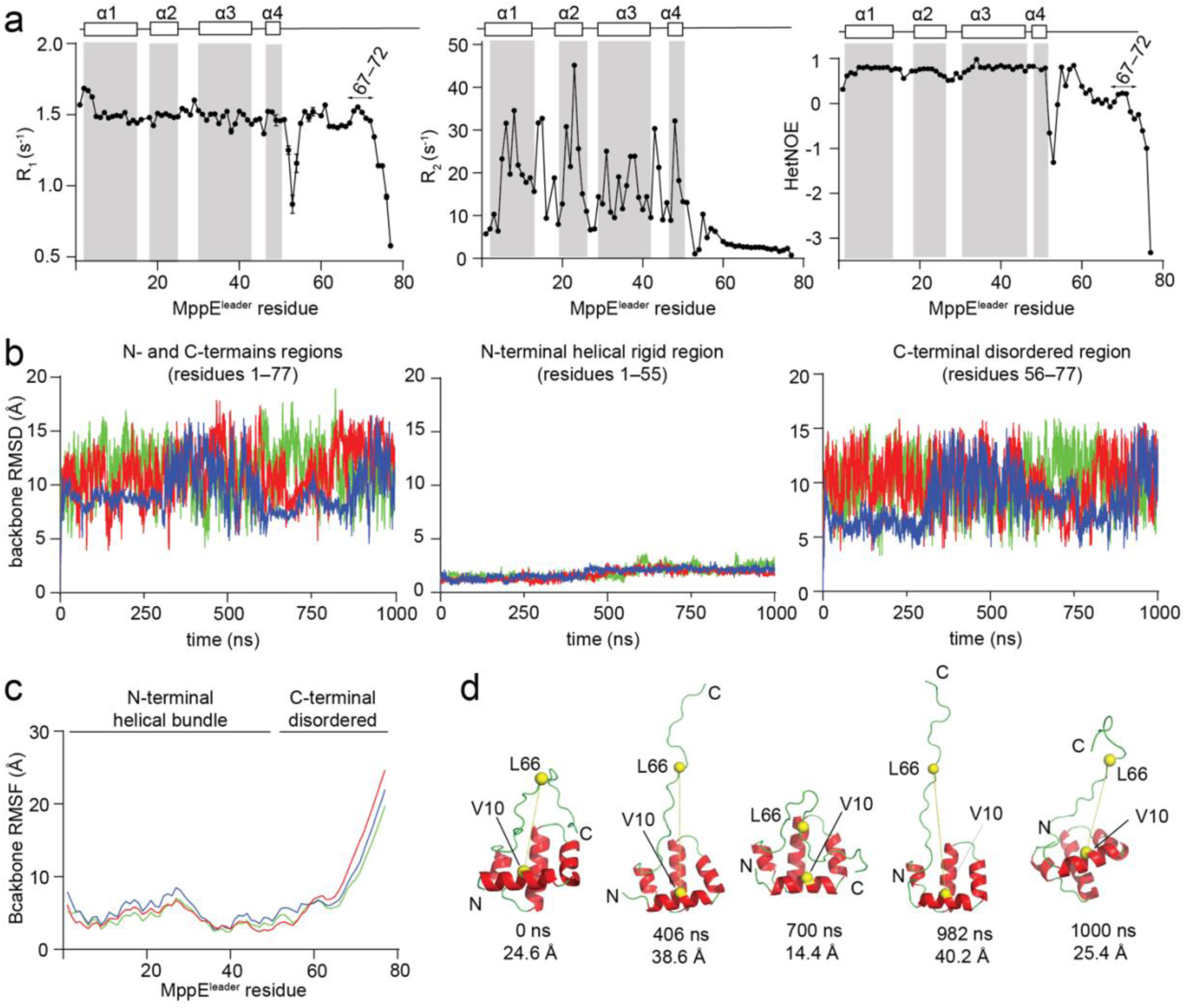
Dynamic properties of the Nif11-LP MppE^leader^. **a** Backbone dynamics of the MppE^leader^ obtained from R_1_ (s^−1^), R_2_ (s^−1^), and ^15^N-{^1^H} HetNOE measurements recorded at 25°C at a ^1^H field of 800 MHz. A cartoon schematic of the MppE^leader^ secondary structure is provided for reference. Gray boxes highlight structured regions. Error bars for the R_1_ and R_2_ values were determined using a statistical bootstrapping scheme in NMRFAM-Sparky. Errors bars for hetNOE values were determined from spectral noise in NMRFAM-Sparky. Residues 67–72 that possess slightly higher R_1_ and HetNOE values are indicated. **b** Plot of backbone RMSD versus time for MppE^leader^ calculated relative to the input structure along three independent 1,000 ns trajectories. For comparison, RMSD calculations were performed on MppE^leader^ separately for left: combined N- and C-terminal regions (residues 1–77), middle: the N-terminal region (residues 1–55), and right: the C-terminal region (residues 56–77). **c** Plot of backbone RMSF versus MppE^leader^ residue number along three independent 1,000 ns trajectories. **d** Representative structural snapshots of MppE^leader^ from MD simulations at specific timepoints with distance measurements between the V10 Cα atom (located on α1 helix) and L66 Cα atom (located on the C-terminal L(X)_4_L motif).

To gain further insight into the dynamic nature of Nif11-LPs, we investigated the backbone dynamics of MppE^leader^ using all-atom molecular dynamics (MD) simulations performed in explicit solvent across three independent 1,000 nanosecond trajectories. Analysis of the backbone root mean square deviation (RMSD) and root mean square fluctuation (RMSF) revealed rigidity within the structured N-terminal domain and substantial conformational flexibility in the C-terminal region of the MppE^leader^ (Fig. 5b–5c, Fig. S22). A GROMACS cluster analysis revealed large-scale movement of the C-terminal residues, while α1–α4 remained rigidly in place (Fig. 5d). Mapping the C-terminal dynamics and its contribution to the overall RMSDs was enabled by monitoring the time-dependent change in the distance between the Cα atoms of Val10 (located on α1) and Leu66 (Fig. 5d). These data support prior inferences that MppE^leader^ exhibits a rigid N-terminal tetrahelical core and a highly dynamic C-terminal region.

### Mapping interactions of MppE^leader^ with MppG

The lack of production of mono- or dihydroxylated products in the absence of the MppE^leader^ allows us to posit that MppG would recognize and bind to MppE^leader^ (Fig. 2e–2f). To discern the molecular details for MppE^leader^ engagement with MppG, we acquired 2D ^1^H-^15^N SOFAST HMQC spectra for U-[^15^N]-labeled MppE^leader^ in the absence and presence of increasing molar ratio of unlabeled MppG (Table S5). Even at high stoichiometric ratio of MppG, MppE^leader^ demonstrated high-quality NMR spectra denoting preservation of the Nif11-related structure upon interaction with MppG (in blue in Fig. 6a, Fig. S23–S27). Association with MppG induced major chemical shift perturbations in MppE^leader^. Instability of MppG precluded acquisition of 3D spectra for assignment of the bound complex. Despite this limitation, with only 2D data in hand, several amino acid peaks could readily be annotated as shifting upon interaction with MppG. Among these were backbone amide cross peaks corresponding to the MppE^leader^ residues Ile6, Phe9, Ile13, Leu19, Leu23, Val36, and Ile39 (Fig. 6b). As noted above, these residues are highly conserved in Nif11-LPs and their side chains construct the hydrophobic nucleus around which the Nif11-LP tetrahelical bundle is organized. The observation that the backbone amide cross peaks for these residues were perturbed upon titration with MppG implies that the rigidly structured tetrahelical bundle of MppE^leader^ makes intimate contact with MppG.

**Fig. 6.**
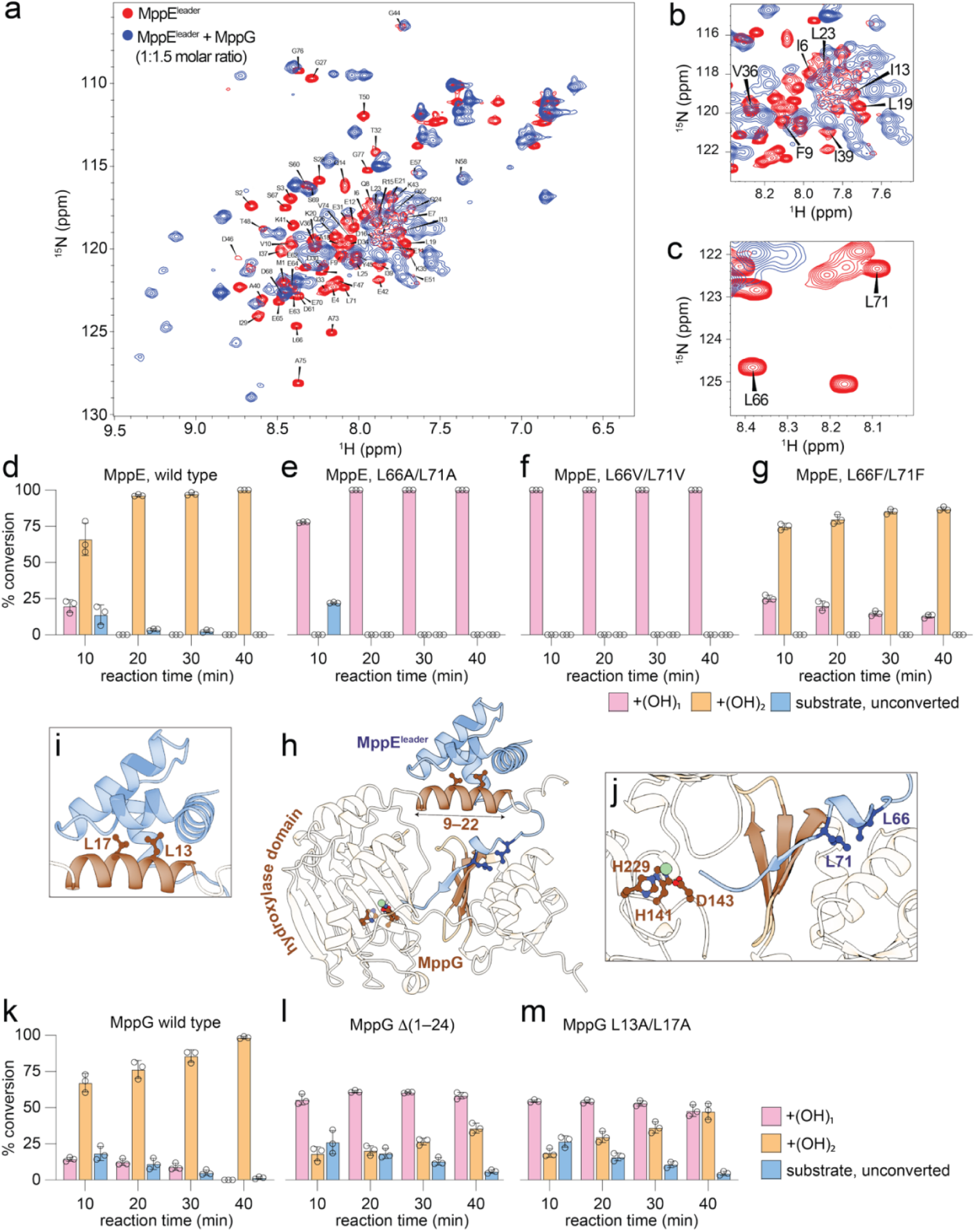
Interaction of Nif11-LP MppE^leader^ with MppG. **a** 2D ^1^H-^15^N SOFAST HMQC spectra of 100 μM U-[^15^N,^13^C]-labeled MppE^leader^ peptide in the absence (in red) and presence of 1.5 molar excess of unlabeled MppG (in blue) acquired at a ^1^H field of 800 MHz at 25°C. **b** Zoomed in view of the overlaid spectra with the backbone amide cross peaks for residues that construct the hydrophobic nucleus in the MppE^leader^ N-terminal tetrahelical bundle labeled. **c** Zoomed in view of the overlaid spectra with the backbone amide cross peaks for MppE^leader^ Leu66 and Leu71 residues labeled. These two residues constitute the L(X)_4_L motif in MppE. **d** Time-dependent conversion of wild-type MppE in which the MppE^core^ was macrocyclized by MppM. Denoted are relative abundances of the mono- and di-hydroxylated products (pink and brown, respectively) and of the unconverted substrate (blue). Time-dependent product formation when the Leu66 and Leu71 residues that constitute the L(X)_4_L motif in MppE were mutated to **e** Ala, **f** Val, and **g** Phe. **h** AlphaFold 3-predicted model of MppE^leader^ (in blue) binding to MppG (brown). The side chains MppG residues His141, Asp143, and His229 that coordinate the Fe-ion (in green, sphere) in the MppG hydroxylase domain are shown in stick-ball representation. The antiparallel three stranded β-sheet in the MppG RRE domain is colored darker for emphasis. Also colored darker is a *a*-helix comprising of MppG residues 9–22 that coordinates to the MppE^leader^ tetrahelical bundle. **i** Zoomed in view of the side chains of MppG residues Leu13 and Leu17 that are modeled to be inserted into the hydrophobic nucleus of MppE^leader^. **j** Zoomed in view of extension of the MppG RRE domain’s antiparallel β-sheet by an additional β-strand contributed by the C-terminal region of MppE^leader^. The side chains of MppE residues Leu66 and Leu71 are shown in stick-ball representation. Note that in the AlphaFold 3-generated model, the MppE^leader^ terminates proximal to the MppG active site as denoted by the site of Fe-coordination. Time-dependent mono- and di-hydroxylated product formation, and leftover unconverted substrate when wild type MppE with macrocyclic MppE^core^ was provided to **k** wild type MppG, **l** MppG in which residues 1–24 were deleted, and **m** MppG wherein residues Leu13 and Leu17 were mutated to Ala.

Also conspicuous among the cross peaks that were perturbed upon titration with MppG were those corresponding to the backbone amides for MppE^leader^ residues Leu66 and Leu71 (Fig. 6c). These two Leu residues constitute the L(X)_4_L motif (Fig. 4c). The L(X)_4_L motif is positioned distal from the tetrahelical bundle in the unstructured loop region spanning MppE^leader^ residues 52–77. The L(X)_4_L motif is highly conserved in Nif11-LPs and NHLPs. The L(X)_4_L motif is also conserved in other smaller, unstructured RiPP precursor leader peptides such as in the precursor peptides that are morphed into cyanobactins.^19,53^ The leader peptide region containing the L(X)_4_L motif has been experimentally demonstrated to interact with RRE domains in RiPP biosynthetic enzymes such that it extends the three stranded RRE antiparallel β-sheet by forming an additional antiparallel β-strand.^22,23,29^ It should be noted that MppG also possesses an RRE domain at the C-terminus (Fig. S1). RRE-independent modalities that involve hydrophobic interactions mediated by the Leu side chains of the L(X)_4_L motif for binding to RiPP biosynthetic enzymes have also been reported.^21,27^

With the *in vitro* activity of MppG established, we biochemically tested the role of Leu66 and Leu71 side chains in mediating MppG/MppE^leader^ interactions. Expectedly, the wild-type MppE^leader^ supported robust dihydroxylation activity of MppG such that negligible monohydroxylated products or unconverted substrate were detected within 20 min of reaction initiation (Fig. 6d, Fig. S28–S31). However, when the Leu66 and Leu71 residues were mutated to Ala or Val, we noticed that the MppG-catalyzed reactions stalled to generate only the monohydroxylated product; no dihydroxylated product formation was detected (Fig. 6e–6f, Fig. S32–S39). Mutation to Phe restored dihydroxylated product formation, albeit with a small compromise in reaction velocity (Fig. 6g, Fig. S40–S43). Complementing the NMR titration observations, these biochemical data allow us to posit that the Leu66 and Leu71 residues also make intimate contact with MppG. Considering literature precedent, these contacts could involve the MppG RRE domain. Taken together, it was abundantly clear that the N-terminal structured tetrahelical bundle as well as the C-terminal unstructured loop in MppE^leader^ both mediate contact with MppG. This bipartite interaction model is operative for NHLP binding to YcaO cyclodehydratases, and for Nif11-LP and NHLP binding to lanthionine synthetases.^28,29^

### Model of the MppE^leader^/MppG complex

We next sought to generate a structural model of MppG/MppE^leader^ complex that would conform to the abovementioned biophysical and biochemical observations. AlphaFold 3 predicted a MppG/MppE^leader^ complex structure with modest statistics (ipTM=0.5; pTM=0.63) (Fig. 6h).^54^ The structures of the MppG hydroxylase and the RRE domains were modeled with high confidence and resemble experimentally established structures, as was also the case for the MppE^leader^ which was modeled with excellent conformity to the solution structure of the tetrahelical bundle reported above. The Fe ion was modeled to be coordinated by the MppG His141, Asp143, and His229 side chains. Regions in the complex that were modeled with low confidence include the interface between the MppG hydroxylase and RRE domains, the long, 25 amino acid extension at the N-terminus of MppG, and the C-terminal loop region of MppE^leader^ comprising of residues 52–77 (Fig. S44). Despite these shortcomings, there were key observations that could be made which corroborate the NMR chemical shift perturbation and biochemical data presented above.

Firstly, the MppE^leader^ tetrahelical bundle was closely associated with an α-helix at the N-terminus of MppG comprising of residues 9–22 (Fig. 6h). This MppG helix, which makes no discernable contact with either the RRE or the hydroxylase domains of MppG, positions the MppG Leu13 and Leu17 side chains into the hydrophobic nucleus of the MppE^leader^ tetrahelical bundle (Fig. 6i). Secondly, the MppE^leader^ terminated proximal to the MppG active site, as defined by the Fe-ion binding site (Fig. 6j). Here, a part of the C-terminal unstructured loop region of the MppE^leader^ was morphed into a β-strand that extended the MppG RRE domain antiparallel β-sheet in a manner that has been observed in the crystal structures of leader peptide binding to RRE domains of YcaO cyclodehydratases and lanthipeptide dehydratases.^22,23^ The MppE^leader^ Leu66 and Leu71 side chains were buried in a hydrophobic patch on the MppE RRE domain surface. This modeled hydrophobic engagement with the RRE domain is in line with the biochemical data presented above wherein mutation of the MppE^leader^ Leu66 and Leu71 residues disrupts the dihydroxylation activity of MppG (Fig. 6d–6g).

To validate the modeled interaction of the MppE^leader^ tetrahelical bundle with the MppG 9–22 α-helix, we generated a MppG Δ(1–24) variant in which the first 24 amino acids were deleted. The MppG Δ(1–24) variant was purified with no alteration in enzyme oligomeric state as discerned by size-exclusion chromatography (Fig. S45–S46). However, as compared to wild-type MppG, the MppG Δ(1–24) variant demonstrated compromised ability to generate dihydroxylated products (Fig. 6k–6l, Fig. S47–S54). When only the MppG Leu13 and Leu17 residues were mutated to Ala, we observed a similar reduction in substrate turnover to dihydroxylated products (Fig. 6m, Fig. S55–S59). These data support the AlphaFold 3-predicted model and corroborate the experimental observation of chemical shift perturbations such that the hydrophobic nucleus of the MppE^leader^ is intimately engaged with MppG *a*-helix formed by residues 9–22.

It is curious to observe that mutations that disrupt engagement of the C-terminal disordered region MppE^leader^ with the MppG RRE domain stall the reaction at the monohydroxylated intermediate and only retard the production of the dihydroxylated product (MppE Leu66Ala/Leu71Ala and MppE Leu66Val/Leu71Val, Fig. 6e–6f). However, mutations that disrupt the engagement of the N-terminal tetrahelical bundle of MppE^leader^ with MppG retard the mono-, as well as the dihydroxylation events (MppG Δ1–24 and MppG Leu13Ala/Leu17Ala, Fig. 6l–6m). MppG is a processive enzyme catalyzing two modifications upon the MppE^core^. In line with other processive enzymes in RiPP biosynthesis such as lanthionine synthetases, it is plausible that the dynamic reordering of the enzyme/substrate complexes occurs after each catalytic event such that different sets of intermolecular interactions dictate binding of the substrate and of the intermediate(s) to the enzyme.^55^ For instance, engagement of the L(X)_4_L motif of the MppE^leader^ to MppG is principally important to drive the second hydroxylation event, but not the first. In contrast, engagement of the tetrahelical bundle of MppE^leader^ to MppG is important for both hydroxylation events. It is entirely plausible that the macrocyclic MppE^core^ (substrate) and the macrocyclic *and* monohydroxylated MppE^core^ (intermediate) bind to the MppG active site differently—indeed that has to be the case as the two sites of hydroxylation are separated by nine atoms (Fig. 2g). Here, fidelity of interactions of the MppE^leader^ L(X)_4_L motif would be crucial to set the register for proper binding of the macrocyclic, monohydroxylated MppE^core^ to the enzyme active site such that the second hydroxylation event can proceed. Substrate-exercised control in the reactivity of RiPP biosynthetic enzymes has been appreciated in other systems as well.^56–58^ In the absence of the macrocyclic MppE^core^, the model presented in Fig. 6h likely represents an overall “interaction ensemble” of the interaction of MppE^leader^ with MppG without capturing the intricacies of how these interactions would change with alterations in the chemistry of MppE^core^.

The bipartite structures of NHLPs and Nif11-LPs present a rich tapestry of intermolecular interactions to engage RiPP biosynthetic enzymes. The N-terminal structured region of NHLPs, which comprises a trihelical bundle and a two-strand antiparallel β-sheet, mediates contact with lanthionine synthetases and YcaO cyclodehydratases.^27–29^ Among Nif11-LPs, the structured region for MppE^leader^ itself has been shown to engage with the lanthionine synthetase MppM, the prenyltransferase MppC, and the flavin-dependent halogenase MppI encoded in the *mpp* BGC (Fig. 2a).^28,38,39^ Interaction with MppG is the first instance in which the interaction of a Nif11-LP with an RRE has been experimentally validated. It is thus curious to observe that the mode of β-sheet elongation first observed for a short and unstructured cyanobactin leader peptide is conserved by the C-terminal unstructured regions of NHLPs and Nif11-LPs. Findings presented herein illuminate possible structural co-evolution of RiPP precursor peptides and RiPP biosynthetic enzymes. In addition to the RRE domain, MppG possesses a rather isolated N-terminal helical extension to engage with the tetrahelical structured nucleus of the Nif11-LP. These highly specialized interactions are likely to encumber mix-and-match engineering experiments in which enzymes such as MppG may be tasked to modify diverse precursor peptides. Regardless of this possible complication, determining the structures of and inventorying the diverse molecular mechanisms for NHLP and Nif11-LP interaction with RiPP biosynthetic enzymes sets the stage for *de novo* design of non-natural long leader peptide sequences that can interface with RiPP biosynthetic enzymes. The NHFe/α-KG-catalyzed hydroxylation is another PTM that is shared between RiPP biosynthesis and primary biology; the JmjC (Jumonji C) domain-containing NHFe/α-KG hydroxylases modify RNA and proteinogenic substrates as an epigenetic control mechanism.^40,59–61^ Primary and secondary metabolism seems to be picking from a common set of chemical transformations but tailoring the intermolecular substrate/enzyme interaction modalities for varied outcomes.

## Materials and Methods

### General materials and methods

All chemicals, solvents, and media components were obtained commercially from Sigma-Aldrich, Fisher Scientific, and VWR, and used without further purification. Phusion high-fidelity DNA polymerase and Gibson assembly Master Mix were purchased from New England Biolabs. PrimeSTAR DNA polymerase Master Mix was purchased from Takara Bio. Synthetic DNA fragments were ordered from Twist Biosciences. Isotopically labeled U-^15^NH_4_Cl and U-^13^C_6_-labeled glucose were purchased from Cambridge Isotope Laboratories. Mass spectra were recorded on an Agilent 6530 quadrupole time of flight (ToF) mass spectrometer equipped with an electrospray ionization (ESI) source coupled to an Agilent 1260 liquid chromatography system equipped with a diode array detector (DAD). CD spectra were acquired using a Jasco J-815 spectropolarimeter equipped with a circulating water bath.

### Genome mining and generation of a sequence similarity network of Nif11-LP

Sequences of Nif11-LPs were obtained from UniProt v2026_02. Duplicate sequences were removed using seqkit ^62^. The SSN illustrated in Figure 1 was generated from 2,395 FASTA-formatted Nif11-LP sequences using the Enzyme Function Initiative Enzyme Similarity Tool (EFI-EST) with an alignment score threshold of 18 ^63^. SSNs were visualized in Cytoscape v3.10.3 where nodes were colored by bacterial phyla ^64^.

## Preparation of recombinant plasmids

The synthetic gene codon optimized for expression in *E. coli* and encoding MppG was inserted into the pET-28a(+) plasmid to afford a thrombin cleavable N-terminal (His)_6_ tag. The pET-28a(+)-*mppG-L13A/L17A* and pET-28a(+)-*mppG-*(Δ1–24) plasmids were prepared from wild type pET-28a(+)-*mppG* by site-directed mutagenesis. All plasmids were validated by sequencing. Preparation of plasmids that encode the MppE precursor peptide and its variants has been reported elsewhere ^38,39^. See Table S1 for protein sequences used in this study.

## General protein expression

Plasmids for the expression of *mppG* and variants thereof were transformed into *E. coli* BL21 (DE3) under appropriate antibiotic pressure for protein expression. Starter cultures were grown by inoculating 10 mL of terrific broth (TB) containing appropriate antibiotics with single colonies, then shaking the liquid culture at 180 rpm for 18 h at 37 °C. 1 L of autoclaved TB media containing appropriate concentrations of antibiotics was then inoculated with the 10 mL starter culture, followed by incubation at 37 °C while shaking at 180 rpm. When optical density at 600 nm (OD_600_) reached 0.6–0.8, cultures was cooled to 18 °C by adjusting the incubator temperature, and protein expression was induced by addition of 0.15 mM isopropyl-β-D-thiogalactopyranoside (IPTG). The culture was then incubated at 18 °C for 18 h while shaking at 180 rpm. Expression of *mppM*, *mppE*, and variants thereof has been reported previously, and identical procedures were used for this study.^38,39^

## General protein purification

Abovementioned protein expression cultures were harvested by centrifugation (3,300×g, 4 °C, 25 min). Cells obtained from a 1 L culture were resuspended in 40 mL of buffer containing 20 mM Tris-Cl (pH 7.5), 500 mM NaCl, and 30 mM imidazole. Cells were lysed by sonication using a QSonica Q500 (50% amplitude, 10 s on and 50 s off pulses, for 15 min on ice). The supernatant was clarified by centrifugation (39,000×g, 4 °C, 45 min) and the supernatant collected.

Affinity chromatography using Ni-NTA: The following procedure was performed at 4 °C. A 5 mL Ni^2+^-charged column (HisTrap, Cytiva) was equilibrated with 50 mL buffer containing 20 mM Tris-Cl (pH 7.5), 500 mM NaCl, and 30 mM imidazole (binding buffer). The column was then loaded with the clarified culture supernatant at a flow rate of 2 mL/min using an ÄKTA Go FPLC system (Cytiva). The column was washed extensively with the binding buffer, followed by an additional wash with 6 column volumes of 5% buffer containing 20 mM Tris-Cl (pH 7.5), 500 mM NaCl, and 1 M imidazole (elution buffer). Proteins were eluted using 12 column volumes of buffer, with the elution buffer increasing linearly from 5 to 100%. Fractions were analyzed for purity using SDS-PAGE. Fractions containing protein of interest were then pooled and concentrated using centrifugal filters, followed by buffer exchange into a storage buffer containing 20 mM HEPES-Na (pH 7.5), 500 mM NaCl, and 10% glycerol, using a PD-10 column (Cytiva). Protein concentrations were measured using the Bradford assay followed by freezing protein in small aliquots at −80 °C.

Size-exclusion chromatography: Purified fractions of MppG wild-type, MppG-L13A/L17A, and MppG-(Δ1–24) were analyzed for their oligomerization state using a size-exclusion chromatography column (Superdex 200 increase 10/300 GL, Cytiva). The column was calibrated using proteins ranging from 43,000 to 669,000 Da (high-molecular-weight gel filtration calibration kit, Cytiva). The column was equilibrated at a flow rate of 0.3 mL/min in buffer containing 40 mM sodium phosphate (pH 7.5) and 100 mM NaCl. 300 μL of concentrated MppG protein and variants thereof were injected on to the column followed by isocratic elution.

## Expression and purification of isotopically labeled MppE^leader^

Plasmid encoding N-(His)6-MppE^leader^ was transformed into *E. coli* BL21(DE3) under antibiotic selection pressure. The preparation of minimal media proceeded using procedures described previously.^27,28^ 1 L M9 minimal media was supplemented with kanamycin, 3 g of ^13^C-labeled glucose, 1 g of ^15^N-labeled ammonium chloride, and 0.2 g ISOGRO-^13^C,^15^N was inoculated with 5 mL overnight culture. The cultures were incubated at 37 °C with shaking until the OD_600_ reached 0.9. The culture was then cooled down to 18 °C for 1 h before protein expression was induced by adding 0.8 mM IPTG. Induced cultures were further incubated at 18 °C, 180 rpm shaking for 18 h. The cultures were harvested by centrifugation at 3,300×g for 30 min at 4 °C. The collected cell pellet was resuspended in lysis buffer A (20 mM Tris-Cl (pH 7.5), 1 M NaCl) and lysed by sonication as described above. Lysate was removed by centrifugation at 35,000×g for 45 min and the clarified supernatant was collected. The supernatant was loaded on to a 5 mL His-Trap column charged with Ni^2+^ and preequilibrated in lysis buffer. The column was washed with wash buffer B (20 mM Tris-Cl (pH 7.5), 1 M NaCl, 30 mM imidazole) and eluted using a linear gradient from 0% to 100% across 30 min with elution buffer B (20 mM Tris-Cl (pH 7.5), 1 M NaCl, 1 M imidazole). The purified protein was dialyzed overnight at 4 °C against buffer containing 20 mM Tris-Cl (pH 7.5), 100 mM NaCl. The dialyzed protein was concentrated using a 3 kDa Amicon centrifugal filter. Concentrated protein fractions were further purified by size exclusion chromatography using a Superdex 75 16/200 column with buffer containing 20 mM Na-phosphate (pH 6.5), 100 mM NaCl. Fraction containing pure proteins, as judged by SDS-PAGE were pooled and concentrated using a 3 kDa Amicon centrifugal filter at 4 °C.

## Enzymatic assays

All assays were conducted in triplicate, unless otherwise noted. For the endpoint assays to determine a competent substrate for MppG, 100 μL reaction mixtures containing 100 μM linear MppE or MppM-modified MppE, 20 mM HEPES-Na (pH 7.5), 5 mM DTT, 1 mM α-KG, 0.5 mM L-ascorbic acid, 0.1 mM Fe(NH_4_)_2_(SO_4_)_2_, and 10 μM MppG were incubated for 2 h at 30 °C. The reactions were quenched with the addition of 0.025 M HCl (final concentration), then incubated on ice for 5 min, followed by the addition of 0.025 M NaOH. Precipitates were removed by centrifugation (16,800×g, 5 min), followed by the addition of 10 μM LahT150 protease to the supernatant and incubation for 2 h at 30 °C to excise the modified MppE^core^ from the MppE^leader^.^21^ An equal volume of 98% (v/v) MeOH + 2% (v/v) formic acid was added to every reaction mixture followed by centrifugation at 16,800×g for 30 min before analysis using liquid chromatography/mass spectrometry (LC/MS). Note that all reactions performed using MppM-modified MppE and its mutants as substrates required initial incubation of the corresponding partially modified peptide (obtained from co-expression of *mppE* and mutant thereof with *mppM*) with 4 μM purified MppM, 5 mM MgCl_2_, and 0.5 mM ATP for 1 h at 30 °C to obtain MppM-modified peptide in stoichiometric yield.

For the endpoint assay to evaluate the activity of MppG when MppE^leader^ was present *in trans*, 25 μM LahT150 protease was added to 100 μM MppM-modified MppE in a 100 μL reaction containing 20 mM HEPES-Na (pH 7.5) and 5 mM DTT to release the macrocyclized MppE^core^ from MppE^leader21^. The reactions were incubated for 4 h at 30 °C, followed by the addition of 1 mM α-KG, 0.5 mM L-ascorbic acid, 0.1 mM Fe(NH_4_)_2_(SO_4_)_2_, and 10 μM MppG, and then incubated for 2 h at 30 °C. In the endpoint assay to evaluate the activity of MppG in the absence of MppE^leader^, 25 μM LahT150 protease was added to 100 μM MppM-modified MppE in 100 μL reaction mixture containing 20 mM HEPES-Na (pH 7.5) and 5 mM DTT, followed by incubation for 4 h at 30 °C. MppE^leader^ was then precipitated by the addition of 0.07 M HCl (final concentration), followed by addition of 0.07 M NaOH (final concentration) and centrifugation (16,800×g, 5 min) to obtain a clarified reaction mixture. 1 mM α-KG, 0.5 mM L-ascorbic acid, 0.1 mM Fe(NH_4_)_2_(SO_4_)_2_, and 10 μM MppG were added to the clarified reaction mixture, which was then incubated for 2 h at 30 °C.

For the time-course assays to evaluate the activity of MppG with MppE-L(X)_4_L mutants, 220 μL reaction mixtures containing 100 μM MppM-modified MppE or MppE-L(X)_4_L mutants, 20 mM HEPES-Na (pH 7.5), 5 mM DTT, 1 mM α-KG, 0.5 mM L-ascorbic acid, 0.1 mM Fe(NH_4_)_2_(SO_4_)_2_, and 4 μM MppG were incubated at 30 °C. 50 μL aliquots were withdrawn at 10 min intervals over 40 min. Aliquots were treated with 0.025 M HCl (final concentration), incubated on ice for 5 min, and then treated with 0.025 M NaOH (final concentration). Clarified supernatants were obtained by centrifugation (16,800×g, 5 min), treated with 2 μg/mL Glu-C protease (Promega), and then incubated for 4 h at 30 °C to excise the modified MppE^core^ from the MppE^leader^ prior to LC/MS analysis. Time-course assays to evaluate the activities of MppG mutants were performed under the same conditions, using MppM-modified MppE as the substrate and LahT150 protease to excise the MppE^leader^.

## LC/MS analysis protocol

As described above, MppE^leader^ was removed by proteolysis by LahT150, or Glu-C, and precipitated from solution by addition of an equal volume of 98% (v/v) MeOH + 2% (v/v) formic acid to the reaction mixtures. Precipitates were removed by centrifugation (16,800×g, 30 min). This treatment allowed the modified MppE^core^ to remain in solution amenable for LC/MS analysis. Clear mixtures were analyzed on an end-capped 100 mm long C_18_ reversed-phase column with a pore size of 120 Å, 2.7 μm particle size, and 4.6 mm inner diameter (Agilent), using 98% solvent A (H_2_O + 0.1% (v/v) formic acid) for 5 min, increasing solvent B (MeCN + 0.1% (v/v) formic acid) to 100% in a linear gradient from 5 to 18 min, and using 100% solvent B from 18 to 22 min, at a flow rate of 0.3 mL/min. Solvent A was then ramped to 98% in a linear gradient over 2 min, holding solvent A at 98% for another 6 min. Data acquired by the mass spectrometer were analyzed using Agilent MassHunter Qualitative Analysis software. To evaluate product formation, ions corresponding to substrate (**a**), monohydroxylated product (**b**), and dihydroxylated product (**c**) were extracted. The three species (**a**, **b**, and **c**) were observed as [M+2H]^2+^ and [M+H]^+^ ions, with [M+2H]^2+^ charged state being dominant. For species **b**, the [M+2H]^2+^ ion was accompanied by its monodehydrated species (**b_1_**) as an in-source dehydration (−2H) fragment. For species **c**, the [M+2H]^2+^ ion was accompanied by its monodehydrated fragment (**c_1_**) and didehydrated fragment (**c_2_**) (Fig. S6). Therefore, the sum of the areas under the extracted ion chromatogram (EIC) peaks corresponding to **b** and **b_1_** was used to quantify the relative abundance of the monohydroxylated product, while the sum of the areas under the EIC peaks corresponding to **c**, **c_1_**, and **c_2_** was used to quantify the relative abundance of the dihydroxylated product. The formulae used were:

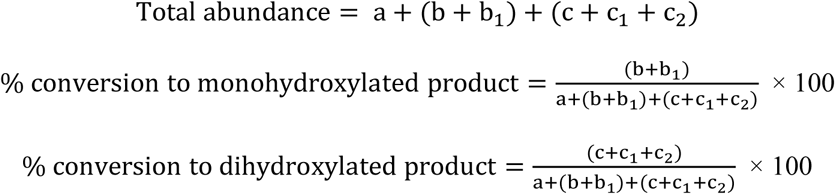

## Production and purification of dihydroxylated MppE^core^ macrocycle for NMR

Gene encoding the N-(His)_6_-MppE-G77K peptide, designed to enable trypsin-mediated excision of the MppE^leader^, was co-expressed with *mppM*. The thusly purified peptide which contained a mixture of linear and macrocyclized MppE-G77K variant was further modified *in vitro* by treatment with MppM to yield macrocyclized MppE-G77K peptide in stoichiometric yield. 350 mg of the MppM-modified peptide was dihydroxylated in batches. Reactions containing 200 μM MppM-modified peptide, 20 mM HEPES-Na (pH 7.5), 5 mM DTT, 2 mM α-KG, 1 mM L-ascorbic acid, 0.2 mM Fe(NH_4_)_2_(SO_4_)_2_, and 40 μM MppG, were incubated for 20 h at 30 °C. The reactions were quenched by treatment with 0.07 M HCl, incubated for 5 min on ice, followed by the addition of 0.07 M NaOH and centrifugation (30,000×g, 4 °C, 30 min). The MppE^leader^ and modified MppE^core^ were separated by treating the resulting clear mixture with trypsin in batches. Reactions contained 50 mM Tris-Cl (pH 8.0), 5 mM DTT, 4 μM trypsin, and ∼160 μM dihydroxylated MppE. These reactions were incubated for 6 h at 30 °C. Following trypsin digestion, reaction mixtures were further treated with 0.1 ng/µL Glu-C, incubated for 6 h at 30 °C. The reaction mixtures were then centrifuged, and the supernatants were loaded on to Strata™-X 33 μm polymeric reversed-phase columns packed with 2 g of sorbent (Phenomenex). Using gravity flow, three Strata™-X 33 μm columns containing 2 g of sorbent were preconditioned with 40 mL MeOH, equilibrated with 40 mL H_2_O, and subsequently loaded with 80 mL of mixture containing soluble peptide fragments from trypsin and Glu-C digestion. The columns were then washed with 160 mL 10% (v/v) MeOH/H_2_O, 40 mL 20% (v/v) MeOH/H_2_O, 120 mL 30% (v/v) MeOH/H_2_O, and 50 mL 40% (v/v) MeOH/H_2_O. Fractions were collected and monitored by LC/MS. The dihydroxylated macrocyclic MppE^core^ typically eluted in 40 mL 60% (v/v) MeOH/H_2_O. MeOH was removed from the pure fractions *in vacuo*, followed by lyophilization to yield ∼20 mg of dihydroxylated macrocyclic MppE^core^ as a white, low-density solid.

After purification on Strata-X column, dried dihydroxylated macrocyclic MppE^core^ was resuspended in HPLC-grade H_2_O and further purified on Agilent 1260 Infinity II HPLC system using a semi-preparative reversed-phase column (Phenomenex Luna, 5 µm C_18_ 100 Å LC column 250×10 mm). The mobile phases did not contain any acid due to the susceptibility of dihydroxylated macrocyclic MppE^core^ to dehydration. The mobile phase solvents used were water (solvent A) and MeCN (solvent B). The elution profile was as follows: 95% of solvent A from 0 to 10 min, followed by a linear gradient increasing solvent B to 23% from 10 to 30 min. Solvent B was then increased to 100% in a linear gradient over 2 min, holding solvent B at 100% for 6 min. Solvent B was then decreased to 5% in a linear gradient over 2 min, holding solvent B for 5 min. Elution fractions were separated by monitoring UV absorbance at 210 and 280 nm. Isolated single fractions containing dihydroxylated macrocyclic MppE^core^ from purification batches were pooled and flash-frozen in dry ice/acetone bath to remove the solvents by freeze-drying.

NMR data were obtained using a Bruker AV-III spectrometer operating at 800.10 MHz and running TopSpin 3.2 software. A conventional 5mm NMR tube was used with a sample volume of ca. 0.5 mL, and data were obtained using a QCI cryoprobe with a sample temperature of 298 K. Standard Bruker pulse programs were used for all experiments. 1D ^1^H data was obtained using presaturation (pulse program zgpr) to reduce the water peak; a 90-degree RF pulse was used, data from 8 scans was retained after 4 dummy scans were completed, with an acquisition time of 2.2 sec and a relaxation delay of 2.0 sec. Data processing used exponential multiplication with a line broadening factor of 1. 1D ^13^C data was acquired using the zgpg30 pulse sequence; 16,000 scans were obtained with an acquisition time of ca. 0.6 sec and a relaxation delay of 2 sec (total time ca. 12 h). Data processing used exponential multiplication with a line broadening factor of 3 followed by baseline straightening. 2D data were acquired and processed as outlined below. Presaturation was used for the homonuclear experiments. The “Experiment” entry also includes any relevant optimization setting. Baseplane correction was applied to all 2D datasets after processing.

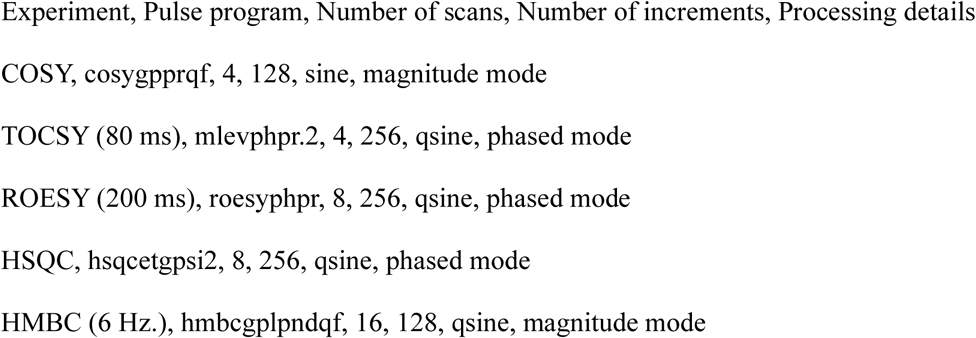

## Circular dichroism spectroscopy (CD)

CD spectroscopy was performed on SEC purified 5 μM MppE^leader^ peptide prepared in 20 mM Na-phosphate (pH 6.5), 20 mM NaCl buffer. Thermal denaturation experiments were performed by increasing the temperature from 10 °C to 90 °C in 1 °C/min increments with acquisition wavelength for CD spectra ranging from 200–300 nm in 0.1 cm path length cuvette. Three replicate spectra, scanned from 200–300 nm at a rate of 500 nm/min, were averaged. Each averaged spectrum was background-corrected and converted to molar circular dichroism [Δε] using the formula

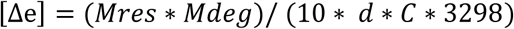

Deconvolution of CD spectra was performed on the BeStSel webserver: https://bestsel.elte.hu/index.php.^48^ Melting temperature was determined by fitting the thermal denaturation curve using a Boltzmann sigmoid equation in GraphPad Prism version 10.

## NMR spectra acquisition, chemical shift assignment, and structure determination

Unless stated otherwise, all solution NMR experiments were conducted at 25 °C in 3 mm NMR tubes in a TCI cryoprobe equipped Bruker Avance III HD 800 MHz spectrometer at the Georgia Tech NMR facility. All data were collected using TopSpin v3.5pl5, processed in NMRPipe v11.5 and analyzed in NMRFAM-SPARKY v3.19.^65,66^

Chemical shifts were assigned after acquiring 3D NMR data on 328 μM U-[^15^N,^13^C]-labeled MppE^leader^ in buffer containing 20 mM Na-phosphate (pH 6.5), 100 mM NaCl, 10% v/v D_2_O. Backbone resonances were assigned using sequential assignment strategies with standard triple-resonance experiments acquired with non-uniform sampling with 18 to 20% Poisson Gap sampling schedule and reconstructed with istHMS.^67^ The following NMR pulse sequences were used: 3D HNCO (Bruker pulse sequence hncogp3d), 3D HNCA (Bruker pulse sequence hncagp3d), 3D HNCACB (Bruker pulse sequence hncacbgp3d), 3D CBCACONH (Bruker pulse sequence cbcaconhgp3d), and 3D amide-amide SOFAST NOESY (pulse sequence sfnoesyNhmqc3d.HnHa-NHn).^68,69^ Acquisition times were set to 79 msec in direct ^1^H, 11 msec in indirect ^15^N, 23 msec in indirect ^13^CO, 10.5 msec in indirect ^13^Cα, and 5 msec in indirect ^13^Cβ. Recycle delay (d1) was set to 1 sec in 3D experiments except for the NOESY. 3D amide-amide SOFAST NOESY experiments were collected with 256 scans, 0.2 sec recycle delay, 350 msec mixing time (d8), and acquisition times set to 79 msec in direct ^1^H, 25 msec in indirect ^15^N, and 16 msec in indirect ^1^H_N_. NOE cross-peaks were assigned manually in NMRFAM-SPARKY.^70^

## Structure calculation in CS-Rosetta

Structure calculations for MppE^leader^ were performed using automated Python scripts with the CS-Rosetta3 toolkit v3.354. Secondary structure index (SSI), random coil index order parameter (RCI-S^2^), and dihedral angle restraints were determined by TALOS-N using assigned chemical shifts of MppE^leader^. The abrelax CS-Rosetta protocol used protein sequence, 3-mer/9-mer fragments, backbone chemical shifts, and NOE restraints obtained from 3D amide-amide SOFAST NOESY experiments. From the 10,000 models calculated the 10 lowest energy models were selected, that fit the experimental NOE data, and dihedral angle restraints. Final structures were validated with MolProbity.^71^

## NMR dynamics measurements for MppE^leader^

U-[^15^N,^13^C]-labeled MppE^leader^ was prepared in NMR buffer containing 10% D_2_O as described above. Longitudinal (T_1_) (Bruker pulse sequence hsqct1etf3gpsi3d) and transverse (T_2_) (Bruker pulse sequence hsqct2etf3gpsi3d) backbone relaxation data were acquired at 25 °C as a series of 2D ^1^H-^15^N HSQC spectra was used to determine backbone ^15^N amide relaxation rates ^52^. The relaxation delay times for T_1_ increments were selected within the range of 10–2,400 msec (10; 20; 60; 100; 200; 400; 600; 800; 1,200; 1,600; 2,000; 2,400; 60; 60), while the values for T_2_ increments were selected between 16.96 to 237.44 msec (16.96; 33.92; 67.84; 101.76; 135.68; 169.6; 203.53; 237.44; 33.92; 33.92). Recycle delay (d1) was set to 3 sec for T_1_ and 1.5 sec for T_2_. The heteronuclear ^15^N{^1^H}-NOE (Bruker pulse sequence hsqcnoef3gpsi3d) was measured by acquiring two 2D ^1^H-^15^N HSQC type spectra with and without proton saturation times of 3 sec. The heteronuclear NOE values were determined from the ratio of intensities of peaks in the two spectra (saturated/unsaturated). Recycle delay (d1) was set to 3 sec for the hetNOE experiment. The relaxation rates (R_1_ = 1/T_1_ and R_2_ = 1/T_2_) were calculated by fitting the signal decay rates to an exponential decay model in NMRFAM-SPARKY.^52^

## NMR titrations

Six different samples each containing U-[^15^N]-labeled 100 μM MppE^leader^ with increasing concentrations of MppG with [MppE]:[MppG] molar ratios of 1:0, 1:0.25, 1:0.75, 1:1, 1:1.5 and 1:2.5 were prepared. Using these samples, 2D ^1^H-^15^N amide SOFAST-HMQC spectra (pulse sequence IBS_SOFAST.x) were acquired with 50% non-uniform sampling (NUS) using the Poisson Gap sampling method with 256 scans and 0.2 s recycle delay with 79 ms and 63 ms acquisition times in the direct ^1^H and indirect ^15^N dimension, respectively, on all six samples. NUS schedules were prepared using the nus@HMS generator v3.^67^

## Molecular dynamics (MD) simulations

All-atom MD simulations in explicit solvent were carried out in GROMACS version 2023.1 using the CHARMM36m force field and TIP3P water model.^72^ CHARMM-GUI solution builder was used to setup simulations.^73^ An integration time step of 2 fs was used with coordinates output every 10 ps. The LINCS algorithm was used to constrain H-bonds. The thermodynamic ensemble was nPT, where the temperature was kept constant at 298.15 K. Nosé–Hoover coupling method was used to maintain temperature with a tau-t of 1 ps. Parrinello–Rahman method was used for an isotropic pressure coupling, with a tau-p of 5 ps and a compressibility of 4.5 × 10−5 bar−1 was used. Short range interactions were treated with a Verlet cutoff scheme with 10 Å electrostatic and van der Waals cutoffs and long-range electrostatics were treated using the particle mesh Ewald method. Periodic rectangular boundaries were used. For MD simulations of MppE, the lowest energy NMR structure (PDB ID 9ZN4) was used. Three independent trajectories (differential initial velocities) for each molecule were acquired for 1 μsec. Simulations were analyzed with VMD v1.9.^74^

## SAXS data acquisition and processing

SAXS experiments were conducted at the Cornell High Energy Synchrotron Source at beamline 7A1 using an x-ray energy of 11.322 keV. The X-ray beam was defined by a slit aperture of 250 µm by 250 µm and attenuated to 5×10^10^‒10^11^ ph/s at the sample. Samples were centrifuged at 13,000 rpm for 10 min at 4 °C prior to measurements to remove any precipitated peptides. A 300 µM MppE^leader^ sample was loaded into the flow cell, continuously oscillated, and measured for 10 to 15 sec, with 1 sec acquisition per data frame. Damage and aggregation were assessed at different beam fluxes. The matching buffer was 20 mM sodium phosphate pH 7.5, 100 mM NaCl. Data were reduced, averaged, and buffer subtracted using BioXTAS RAW.^75^ The scattering profile was fit to one of the top ten NMR-derived structures using FoXS.^76^ GNOM was used to calculate the pair distance distribution function and Dmax.^77^ The electron density was calculated using DENSS, with 100 reconstructions and an oversampling of 5,^78^ and visualized by PyMOL 2.6.2 and ChimeraX.^79^

## Data Availability

Atomic coordinates of the solution NMR structure of MppE^leader^ have been deposited to the Protein Data Bank (PDB) with the accession code 9ZN4. The backbone chemical shift assignments of MppE^leader^ have been deposited to the Biological Magnetic Resonance Bank (BMRB) with the accession code 31287. The SAXS data of MppE^leader^ has been deposited to SASBDB. Molecular dynamics simulations trajectories of MppE^leader^ have been deposited to Zenodo, accessible with the following link: https://doi.org/10.5281/zenodo.20399787.

## Acknowledgements

Authors are thankful for financial support from the National Institutes of Health (NIH) (R35GM142882 to V.A. and R35GM138002 to E.I.P.). This research was supported in part through research cyberinfrastructure resources and services provided by the Partnership for an Advanced Computing Environment (PACE) at the Georgia Institute of Technology, Atlanta, Georgia, USA. The Center for High-Energy X-ray Sciences (CHEXS) at Cornell High Energy Synchrotron Source (CHESS) is supported by the National Science Foundation (NSF) award DMR-2342336, and the MacCHESS resource is supported by NIH award P30GM124166. This study made use of NMRbox: National Center for Biomolecular NMR Data Processing and Analysis, a Biomedical Technology Research Resource (BTRR), which is supported by NIH grant P41GM111135 (NIGMS).

## Ethics Declarations / Competing Interests

The authors declare no competing interests.

